# Sequencing uracil in DNA at single-nucleotide resolution

**DOI:** 10.1101/2021.06.14.448445

**Authors:** Liudan Jiang, Jiayong Yin, Maoxiang Qian, Shaoqin Rong, Kejing Chen, Chengchen Zhao, Yuanqing Tan, Jiayin Guo, Hao Chen, Siyun Gao, Tingting Liu, Yi Liu, Bin Shen, Jian Yang, Yong Zhang, Fei-Long Meng, Jinchuan Hu, Honghui Ma, Yi-Han Chen

**Affiliations:** Department of Cardiology, Shanghai East Hospital, School of Medicine, Tongji University, Shanghai 200092, China; Shanghai Fifth People’s Hospital, Institutes of Biomedical Sciences, Shanghai Key Laboratory of Medical Epigenetics and the International Co-laboratory of Medical Epigenetics and Metabolism, Ministry of Science and Technology, Shanghai Medical College of Fudan University, Shanghai 200032, China; Key Laboratory of Arrhythmias, Ministry of Education, Department of Medical Genetics, School of Medicine, Tongji University, Shanghai 200092, China; Institute of Pediatrics and Department of Hematology and Oncology, Children’s Hospital of Fudan University, the Shanghai Key Laboratory of Medical Epigenetics, Institutes of Biomedical Sciences, Fudan University, Shanghai 200032, China; School of Life Science and Technology, Tongji University, Shanghai, 200092, China; State Key Laboratory of Reproductive Medicine, Nanjing Medical University, Nanjing 211166, China; School of Medicine, Southern University of Science and Technology, Shenzhen 518055, China; State Key Laboratory of Molecular Biology, Shanghai Institute of Biochemistry and Cell Biology, Center for Excellence in Molecular Cell Science, Chinese Academy of Sciences, Shanghai 200031, China; Research Units of Origin and Regulation of Heart Rhythm, Chinese Academy of Medical Sciences, Shanghai 200092, China

## Abstract

As an aberrant base in DNA, uracil is generated by dUMP misincorporation or cytosine deamination, and involved in multiple physiological and pathological processes. Current methods for whole-genome mapping of uracil all rely on uracil-DNA *N*-glycosylase (UNG) and are limited in resolution or specificity. Here, we present a UNG-independent Single-Nucleotide resolution Uracil Sequencing (SNU-seq) method utilizing the UdgX protein which specifically excises the uracil and forms a covalent bond with the resulting deoxyribose. SNU-seq was validated on synthetic DNA and applied to mammalian genomes. We found that the uracil content of pemetrexed-treated cells fluctuated along with DNA replication timing. We also accurately detected uracil introduced through cytosine deamination by the cytosine base editor (nCas9-APOBEC) and verified uracil occurrence in “WRC” motif within Activation-Induced Cytidine Deaminase (AID) hotspot regions in CSR-activated *UNG* ^−/−^ B cells.

---

Uracil is a pyrimidine that possesses similar chemical structure with thymine and can also form base pair with adenine. Because most DNA polymerases are unable to discriminate thymine and uracil, they occasionally incorporate deoxyuridine (dU) instead of deoxythymidine (dT) in DNA, especially when the synthesis of thymidine is disturbed^1–3^. Besides, uracil can be generated by cytosine deamination either due to spontaneous hydrolysis or catalyzed by the AID/APOBEC (apolipoprotein B mRNA editing enzyme, catalytic polypeptide-like) family proteins^3^ and cause C to T substitution if unrepaired. Uracil can be efficiently excised from deoxyribose mainly by UNG^3–4^, generating an apyrimidinic site (AP site) which is further processed by the base excision repair (BER) pathway^5–6^. Although the steady-state frequency of uracil in mammalian genomes is very low (~10^−6^ per nucleotide^7^), it plays crucial roles in diverse biological processes. If uracil is continually incorporated into DNA, hyperactive BER will lead to DNA breaks and even cell death. This so-called “thymine-less cell death”^8,9^ has been exploited in several chemotherapeutic agents. On the other hand, two essential processes during B cell maturation, somatic hypermutation (SHM) and class switch recombination (CSR), are initiated by AID-catalyzed cytosine deamination in immunoglobin genes^10–12^. Furthermore, dysregulation of APOBEC can accumulate undesired C to T mutations in genome, which might facilitate the progression of specific cancer subtypes^13^. Meanwhile, cytosine deamination-mediated C to T conversion has been applied in the genome-editing tool CBE (cytosine base editor) that holds great potential for the treatment of genetic diseases^14–15^. Thus, there is a growing demand for mapping uracil in the whole genome.

Recently, a couple of genome-wide uracil-sequencing methods emerged (reviewed in ref^16^). They generally utilized UNG to convert uridines to AP sites, followed by chemical labeling^17,18^ or incision by an AP endonuclease to generate free 3’OH for further capture^19,20^. Alternatively, an excision-defective UNG-derivate was applied to pull down uracil-containing DNA fragments^21^. Therefore, these assays might be interfered by pre-existing DNA strand breaks, AP sites or other modified nucleotides with aldehyde groups including 5-formyldeoxyuridine (5fU) and 5-formyldeoxycytosine (5fC), although appropriate pre-treatment might reduce these interferences^19^. Moreover, among these methods, only Excision-seq and AI-seq reached single nucleotide resolution^17, 20^. Excision-seq required close uracil on both strands to generate proper double-stranded DNA fragments by UNG / Endo IV treatment^20^. Thus, it has only been applied to genomes with high-density of uracil, e.g. repair-defective *S. cerevisiae* and *E. coli* (~3-8 deoxyuridines per 10^3^ nucleotides^22–23^). AI-seq chemically converted UNG-generated AP sites to azide-cytosine and determined the positions of these modified bases by comparing with input sequences. Therefore, in order to detect uracil generated from cytosine deamination, a significant ratio of U:G mismatch should be observed in the input sequence, which required a high ratio (20%) of dU at specific loci and more than 20x coverage of sequencing depth for input samples^17^.

To address these issues, we set to establish an UNG-independent method to map genome-wide distribution of uracil. Recently, it has been reported that a novel DNA glycosylase from *Mycobacterium smegmatis*, named UdgX, could specifically recognize and excise uracil in DNA and form an irreversible covalent bond with the deoxyribose (**Figure 1a**). This UdgX-DNA covalent complex was exceedingly stable even in harsh treatments such as SDS, NaOH and heat^24–26^. UdgX has been successfully used to visualize uracil *in situ*^27,28^. Based on these findings, we develop a method named ‘SNU-seq’ (<u>S</u>ingle-<u>N</u>ucleotide resolution <u>U</u>racil <u>seq</u>uencing) by combining the unique property of UdgX and the Damage-seq strategy^29^ of high-fidelity DNA polymerase stalling at the protein-DNA adducts to pinpoint uracil at single base resolution.

**Figure 1.**
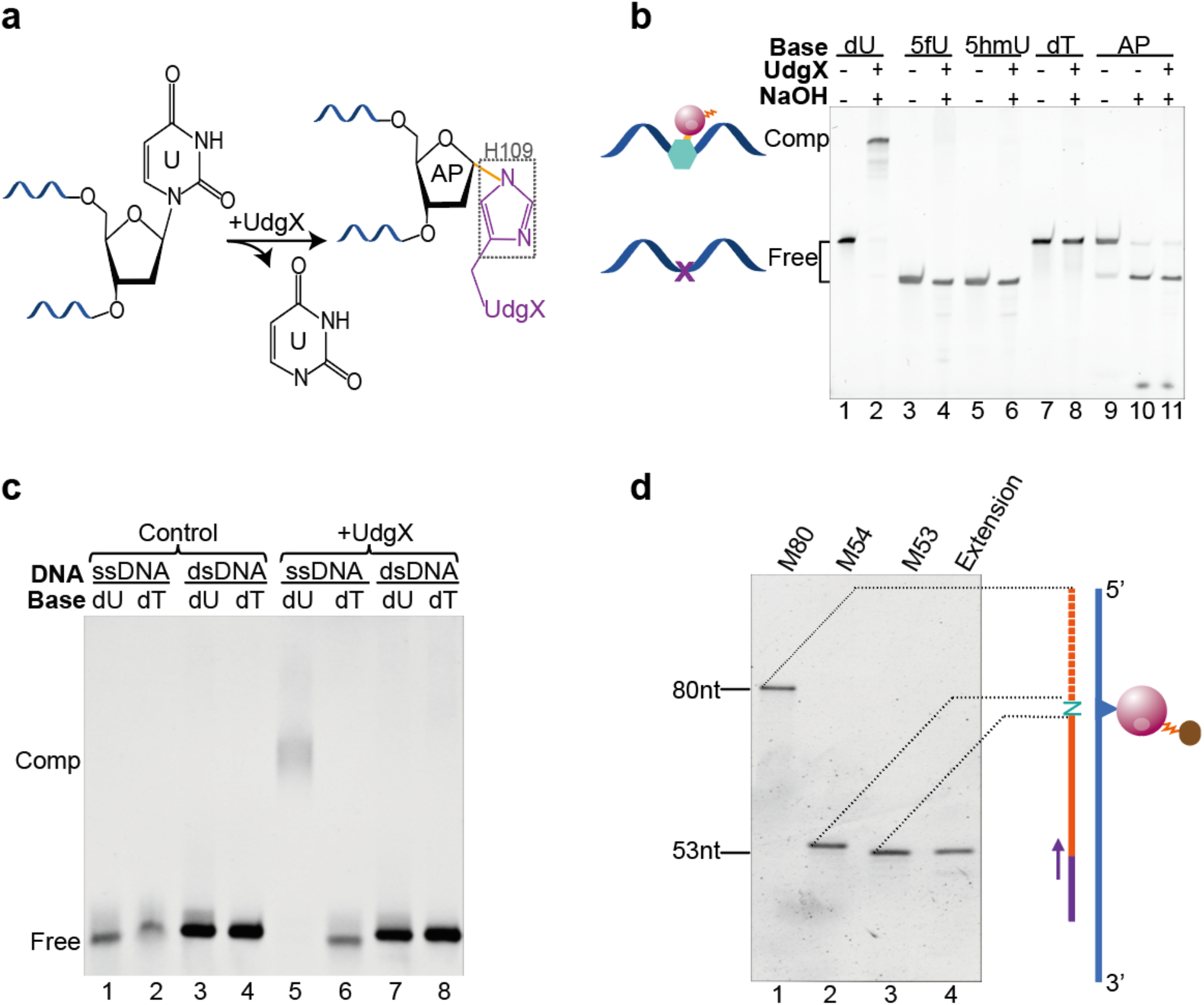
Validation of properties of UdgX. (a) Schematic of UdgX activity to excise uracil and crosslink with resulting AP site by the His109 residue. (b) Reactivity of UdgX with ssDNA harboring various modifications. Reactions were stopped by alkali-heat-treatment which would also cleave AP sites (lanes 10 and 11). (c) Activity of Udgx on ssDNA and dsDNA substrates. (d) Blockage of primer extension by UdgX-dU crosslink. M53, M54 and M80 were markers indicating extension products stopping at one nucleotide before the uracil, at the uracil site, and bypassing the uracil, respectively.

To capture the protein-DNA adducts, a biotin-tag was introduced for its great affinity with streptavidin-beads. To this end, UdgX was fused with N-terminal Avi-tag and co-expressed with BirA ligase which added a biotin to the Avi-tag^30,31^ (**Figure S1, S2a**). As proof of concept, several *in vitro* experiments were performed on synthetic DNA substrates with the biotinylated UdgX. To test the specificity of UdgX to uracil, the protein was incubated with synthetic oligomers harboring dU, dT, AP site and other dT-derivates such as 5hmU and 5fU. The gel shift results showed UdgX quantitatively reacted with uracil only (**Figure 1b**), in line with previous findings^26^. There are conflict reports on the reactivity of UdgX with uracil in double-stranded DNA (dsDNA)^24–25, 27^, our results showed that UdgX could only efficiently crosslink with uracil on single-stranded DNA (ssDNA) but not those on dsDNA (**Figure 1c**), which was consistent with a recent report^28^. We then further optimized the UdgX reaction conditions to maximize the efficiency and minimize possible side reactions (**Figure S2b, S2c**). In Damage-seq, the exact position of targeted damage was detected by the stalling of a high-fidelity DNA polymerase^29^. We therefore performed an extension assay to validate the accuracy of polymerase stalling at UdgX-crosslinked sites. As shown in **Figure 1d**, the extension products (lane 4) were exclusively 53-nt in length, indicating that the polymerase stopped precisely at the nucleotide before the uracil site. Taken together, these results suggested that uracil can be mapped at single-nucleotide resolution by UdgX.

Based on above results, we designed a protocol to detect uracil in genomic DNA. As shown in **Figure 2a**, genomic DNA were sheared, end-repaired and ligated to Adaptor 1. Because UdgX only reacted with ssDNA, the ligated products were denatured before incubating with UdgX. After removing excess free UdgX, the DNA-UdgX complexes were captured by streptavidin-beads. Then a primer was attached and extended by a high-fidelity DNA polymerase. The extension products were released from the beads by alkaline treatment. The eluted DNA were ligated to Adaptor 2 and PCR-amplified to generate sequencing libraries.

**Figure 2.**
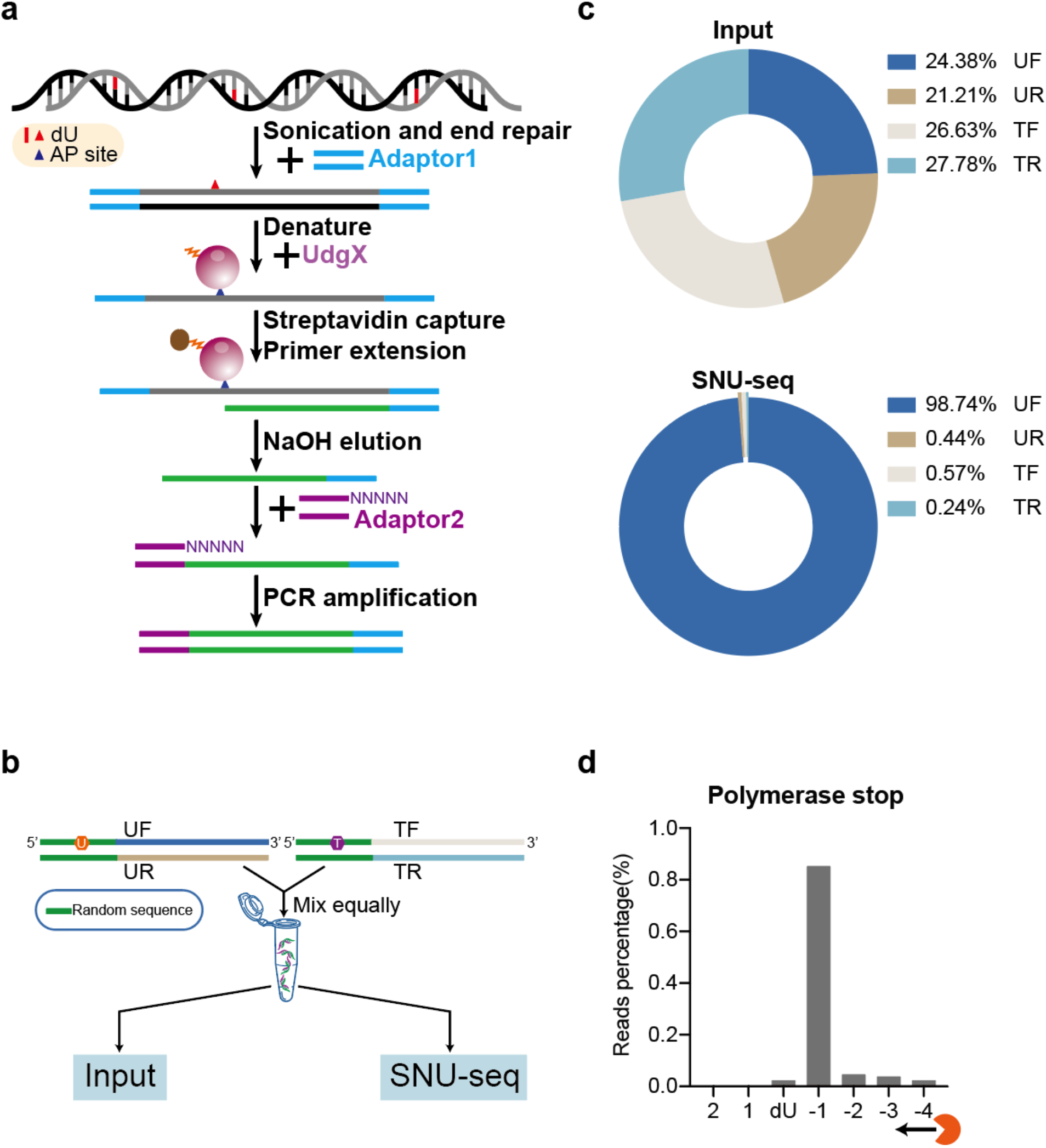
Workflow of SNU-seq and proof of principle with synthetic DNA. (a) Workflow of SNU-seq. (b) Experimental design of input and SNU-seq libraries using synthetic DNA. (c) Distribution of reads aligned to each strand of synthetic DNA. (d) Positions of polymerase stalling on the dU-containing strand.

We first evaluated the SNU-seq workflow with synthetic DNA. Unmodified and dU-containing dsDNA were equally mixed and subjected to input and SNU-seq sequencing (**Figure 2b**). Compared with input, the dU-containing strands (UF) were highly enriched in SNU-seq (~243-fold enrichment, **Figure 2c**). Moreover, the polymerase mainly ended at one nucleotide before the uracil site (~85.2%) (**Figure 2d**).

We next applied SNU-seq to detect uracil in mammalian genome. Pemetrexed (PMX) is an antifolate chemotherapy agent which inhibits thymidylate synthase, leading to accumulation of cellular dUMP and misincorporation of uracil into DNA (**Figure 3a**)^32^. The elevated uracil content in PMX-treated *UNG*^−/−^ HeLa cells (~ 6 per 10^5^ nucleotides) was verified by a modified UdgX-based dot-blot method (**Figure S3b, S3c**). The level of uracil in non-treated sample was too low to be detected, consistent with a previous report that sole lack of UNG could not induce a significant increase of uracil^23^. Genomic DNA of non-treated and PMX-treated cells were prepared for SNU-seq. Quality check of sequencing libraries revealed a higher yield for the PMX-treated sample (**Figure 3b**), suggesting that uracil-containing DNA was enriched. As described in **Figure 2a**, the DNA polymerase stopped just before the damage site, thus the uracil should be at the nucleotide adjacent to the 5′ end of the reads, which ought to be thymine in the reference genome for PMX-treated cells. Thymine was indeed highly enriched at predicted positions in PMX-treated cells but not in untreated cells (**Figure 3c, S4b**). In contrast, cytosine was slightly enriched in untreated cells (30% in SNU-seq vs 20% in the genome), probably because there was more uracil from cytosine deamination when thymidylate synthesis was undisturbed. As uracil was incorporated during DNA replication upon PMX-treatment, we analyzed the uracil distribution in different replicating domains and found that uracil content was lowest in early replicating domains (ERDs) (**Figure 3d, S4a, S4c**), probably due to the fact that ERDs strongly correlate with open chromatin^33^ which is more accessible to other uracil glycosylases (e.g. SMUG1^34^) in UNG knockout cells.

**Figure 3.**
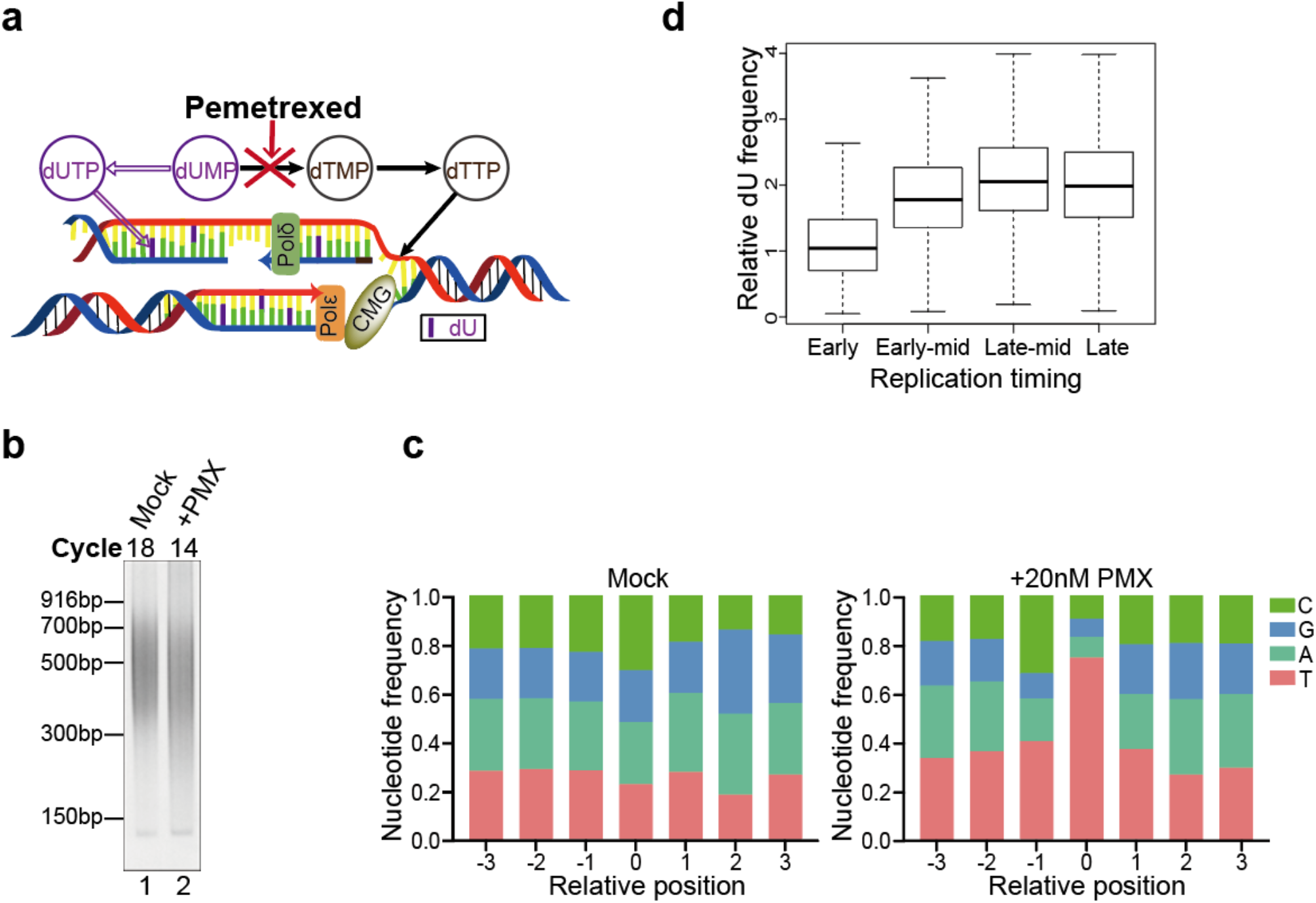
Genome-wide mapping of uracil in pemetrexed-treated cells. (a) Diagram showing that pemetrexed could inhibit dTTP synthesis and lead to misincorporation of uracil into the genome. (b) Quality check of SNU-seq libraries. (c) Nucleotide frequencies around the predicted uracil site. “Position 1” was the first nucleotide of sequencing reads, and “Position 0” was the predicted uracil site. (d) Boxplots showing the frequency of dU in PMX-treated cells as a function of replication timing. Replicate A was shown.

The cytosine base editor (CBE) utilizes a mutated Cas9 and sgRNA to bring a cytosine deaminase (APOBEC) to specific loci and convert C to U on the non-complementary strand, thus achieving C:G to T:A transition (**Figure 4a**)^15,35^. We examined the capability of SNU-seq to detect CBE (AncBE4max)-induced site-specific uracil at *RNF2* loci in HEK293T cells. Sanger sequencing confirmed that the former two of the three cytosines (C_3_, C_6_, C_12_) in the editing window had been effectively edited (**Figure 4b**), consistent with a previous report that AncBE4max showed highest activity at C_6_ and poorest activity at C_12_^35^. Since Sanger sequencing is unable to distinguish dU and dT, we adopted a USER (Uracil-Specific Excision Reagent)-digested LM (ligation-mediated)-PCR assay to measure the occupation of loci-specific uracil (**Figure S5a**, see methods), which revealed that the majority of edited cells contained dU(s) at the *RNF2* loci (**Figure 4c**). As shown in **Figures 4d**and **S5b**, SNU-seq could detect the existence of uracil within the *RNF2* loci, mostly at position C_6_, although C_3_ was also moderately edited. There is a caveat, though, that SNU-seq could only detect the first cytosine from 3’ ends if multiple cytosines were present in the same fragment. To assess the sensitivity of SNU-seq, genomic DNA from the edited sample was diluted with DNA from the unedited sample. It was shown that SNU-seq was reliable for samples diluted in 1:50, and was even able to detect uracil in samples diluted in 1:100 (**Figure 4d, S5b**), implying that SNU-seq was able to detect one edited cell in 50-100 unedited cells.

**Figure 4.**
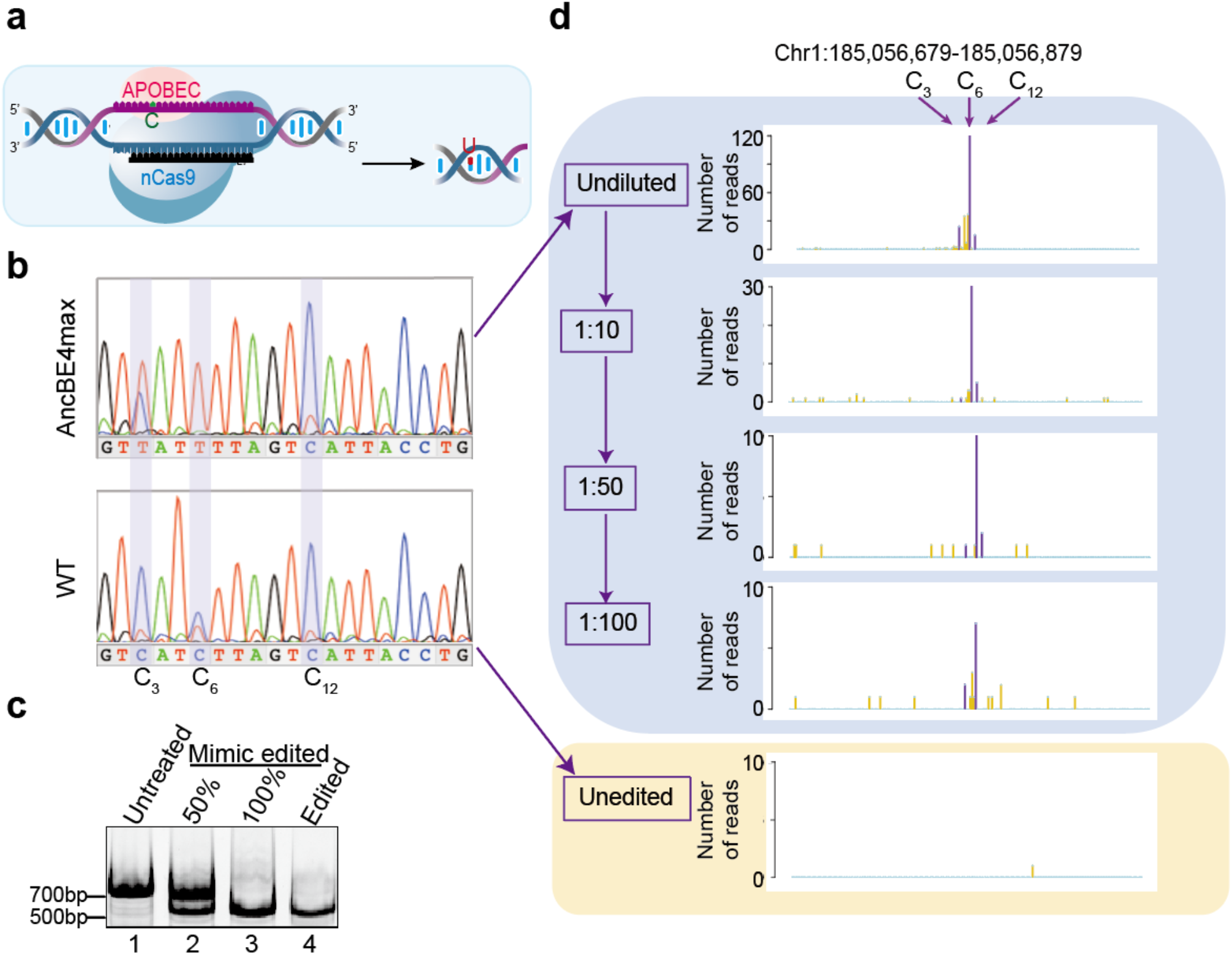
Detection of site-specific uracil by SNU-seq. (a) Schematic of the nCas9-APOBEC-based cytosine base editor. (b) The outcome of gene editing at *RNF2* locus determined by Sanger sequencing. (c) The presence of uracil at *RNF2* loci confirmed by USER-digested-LM-PCR (Figure S5a and methods). NdeI digestion near the editing window was used to mimic USER digestion of dU (lane 3). Untreated and NdeI-digested DNA were mixed equally to simulate 50% C to U conversion (lane 2). (d) Detection of CBE-generated uracil within the editing window by SNU-seq. Edited DNA were diluted with unedited DNA to evaluate the sensitivity of SNU-seq. Replicate A was shown.

The diversity of antibody is dependent on CSR, which is initiated by the incision of dU generated by AID. This enzyme preferentially deaminates cytidines within the 5’ Sμ region and a downstream acceptor S region at WRC (W = A, T; R = A, G) motif, and mostly AGCT motif^36–38^. SNU-seq was used to survey the distribution of uracil in CSR-activated *UNG*^−/−^ B lymphocytes while AID-deficient B lymphocytes were used as a negative control. As shown in **Figure S6**, within the strongest AID-hotspot region^39^, uracil was preferentially located at WRC, especially AGCT motif in *UNG*^−/−^ cells.

In conclusion, we presented an unbiased UNG-independent method, SNU-seq, by which uracil of both origins in the whole genome can be mapped at single nucleotide resolution. Moreover, SNU-seq could easily distinguish the sources of uracil by its original base, thymine or cytosine, in the reference genome. To the best of our knowledge, SNU-seq is the first approach that can directly map site-specific uracil in human genome, even when the ratio of dU is as low as 1% at specific loci. SNU-seq would be valuable for studies on uracil-related questions including the hotspots of dysregulated APOBEC and the efficiency and specificity of CBEs. Notably, since many other proteins can covalently crosslink with specific base-modifications (e.g. HMCES and AP sites ^40^), SNU-seq also provides an innovative strategy for accurate and specific mapping of these base-modification.

## AUTHOR INFORMATION

### Notes

The authors declare no competing financial interest.

### Funding Sources

This work was supported by the National Key Research and Development Plan (2019YFA0801501, to Y.-H.C), the National Natural Science Foundation of China (82088101, 81930013 and 81770397 to Y.-H.C; 31971330 to H.M; 21807013, 31870804 to J.H.), Key Disciplines Group Construction Project of Pudong Health Bureau of Shanghai (PWZxq2017-05), Top-level Clinical Discipline Project of Shanghai Pudong District (PWYgf2018-02), Research Unit of Origin and Regulation of Heart Rhythm, Chinese Academy of Medical Sciences (2019RU045), Innovative research team of high-level local universities in Shanghai and a key laboratory program of the Education Commission of Shanghai Municipality (ZDSYS14005). J.H. is supported by the Program for Professor of Special Appointment (Eastern Scholar) at Shanghai Institutions of Higher Learning, Shanghai Outstanding Young Talent Program, and the innovative research team of high-level local university in Shanghai. M.Q. is supported by the Program for Professor of Special Appointment (Eastern Scholar) at Shanghai Institutions of Higher Learning. Y.-H.C. is a fellow at the Collaborative Innovation Center for Cardiovascular Disease Translational Medicine, Nanjing Medical University.

## Notes

### Competing Interest Statement

The authors have declared no competing interest.

